# Diverse functions of closely homologous actin isoforms are defined by their nucleotide, rather than their amino acid sequence

**DOI:** 10.1101/227546

**Authors:** P. Vedula, S. Kurosaka, N. A. Leu, Y. I. Wolf, S. Shabalina, J. Wang, S. Sterling, D. W. Dong, A. Kashina

**Affiliations:** Department of Biomedical Sciences, University of Pennsylvania, Philadelphia, PA; National Center for Biotechnology Information, National Institutes of Health, Bethesda, MD; Institute for Biomedical Informatics, Perelman School of Medicine, University of Pennsylvania, Philadelphia, PA

## Abstract

β- and γ-cytoplasmic-actin are nearly indistinguishable in their amino acid sequence, but are encoded by different genes that play non-redundant biological roles. The key determinants that drive their functional distinction are unknown. Here we tested the hypothesis that β- and γ-actin functions are defined by their nucleotide, rather than their amino acid sequence, using targeted editing of the mouse genome. Although previous studies have shown that disruption of β-actin gene critically impacts cell migration and mouse embryogenesis, we demonstrate here that generation of a mouse lacking β-actin protein by editing β-actin gene to encode γ-actin protein, and vice versa, does not affect cell migration and/or organism survival. Our data suggest that the essential *in vivo* function of β-actin is provided by the gene sequence independent of the encoded protein isoform. We propose that this regulation constitutes a global “silent code” mechanism that controls the functional diversity of protein isoforms.

## Introduction

Actin is an essential and abundant intracellular protein that plays a major role in developmental morphogenesis, muscle contraction, cell migration, and cellular homeostasis. Two of the closest related actin isoforms, non-muscle β-actin and γ-actin, are ubiquitously expressed but encoded by different genes, producing nearly identical proteins except for 4 residues within their N-termini (Vandekerckhove and Weber, 1978). Notably, β-and γ-actin mRNA coding sequences differ much more significantly, by nearly 13%, due to silent substitutions affecting approximately 40% of their codons (Erba et al., 1986). Recent evidence suggests that this difference in mRNA coding sequence affects translation dynamics of the two actin isoforms: β-actin is translated in bursts and accumulates faster than γ-actin (Buxbaum et al., 2014; Zhang et al., 2010). This difference leads to their differential post-translational modification by arginylation, which targets only β- but not γ-actin (Zhang et al., 2010). Thus, actin isoforms are differentially regulated via changes in their mRNA coding sequence that can affect their translation and post-translationally modified state.

Despite their high similarity and abundance in the same cell types, β- and γ-actin play distinct non-redundant biological roles. A body of evidence shows that these actins localize to different parts of the cell and tend to incorporate into different actin cytoskeletal structures (Dugina et al., 2009; Kashina, 2006; Otey et al., 1986). More definitively, several studies show that β-actin knockout in mice results in early embryonic lethality despite proportional up-regulation of other actin isoforms to compensate for the total actin dosage (Bunnell et al., 2011; Shawlot et al., 1998; Shmerling et al., 2005; Strathdee et al., 2008; Tondeleir et al., 2013; Tondeleir et al., 2014), while γ-actin knockout in mice has a much milder phenotype that does not interfere as significantly with animal survival (Belyantseva et al., 2009; Bunnell and Ervasti, 2010). This lethality of β-actin knockout in mice can be rescued by a targeted knock-in of the β-actin cDNA (Tondeleir et al., 2012), suggesting that the coding region of this gene is far more important for organism’s survival than any non-coding elements affected by the β-actin knockout. The underlying mechanisms conferring such functional differences to these two nearly identical protein isoforms are unknown.

Several distinct features of the two actin isoforms have been proposed as the mechanism conferring unique roles to the two proteins. Some biochemical differences between β- and γ-actin were observed in polymerization assays both *in vitro* and *in vivo* (Bergeron et al., 2010; Kapustina et al., 2016; Muller et al., 2013). In addition, β-actin mRNA, unlike γ-actin mRNA, gets spatially targeted to the cell periphery via zipcode-mediated transport (Hill and Gunning, 1993; Kislauskis et al., 1997). Finally, β- and γ-actin actin can differentially regulate gene expression of a subset of cytoskeleton proteins (Tondeleir et al., 2013; Tondeleir et al., 2014). Despite these differences, no study to date has definitively identified the key functional determinants that confer unique functions to the mammalian actin isoforms in organism’s survival, or determined whether these determinants reside at the amino acid level.

Here we used targeted editing of the mouse genome to test the hypothesis that β- and γ-actin functions *in vivo* are defined by their nucleotide, rather than their amino acid sequence. Although previous studies have shown that disruption of the β-actin gene critically impacts embryonic development and organism survival (Bunnell et al., 2011; Shawlot et al., 1998; Shmerling et al., 2005; Strathdee et al., 2008; Tondeleir et al., 2013; Tondeleir et al., 2014), we demonstrate here that editing of the β-actin coding sequence to encode γ-actin protein without disruption of the rest of the β-actin gene does not affect mouse survival or produce a visible phenotype at the organismal or cellular level. This result shows that γ-actin protein is functionally capable of substituting for β-actin in the absence of gene disruption. Thus, we demonstrate that the differences in *in vivo* functions of β- and γ-actin actin are ultimately determined by their nucleotide rather than amino acid sequence.

## Results

### β-actin nucleotide sequence, rather than amino acid sequence, defines its essential role *in vivo*

It has been previously found that knockout of β-actin gene in mice, unlike γ-actin, leads to early embryonic lethality – a result that definitively demonstrates its essential, non-redundant biological function (Bunnell et al., 2011). To test whether this essential function of β-actin is defined by its amino acid or nucleotide sequence, we used CRISPR/Cas9-mediated gene editing to introduce five point mutations into the native mouse β-actin gene *(Actb)*, altering it to encode γ-actin protein without changing any of the features of the rest of the gene sequence (“beta-coded gamma actin”, Fig. 1A). We termed this edited gene *Actbc-g*, in which the native β-actin gene is nearly intact (with five point mutations within the first 10 codons) and contains the same promoter, as well as the same coding and non-coding elements, but the protein produced from this gene is identical to γ-actin. The outcome is no β-actin protein at all, enabling us to definitively test whether β-actin amino acid sequence, or its nucleotide sequence, is responsible for its essential function in organism’s survival. If intact β-actin amino acid sequence is required, the mutant mice would die in embryogenesis, similarly to the β-actin knockout mice (Bunnell et al., 2011; Shawlot et al., 1998; Shmerling et al., 2005; Strathdee et al., 2008; Tondeleir et al., 2013; Tondeleir et al., 2014). If the nucleotide sequence also contributes, these mice would be expected to survive longer than the β-actin knockout mice and have an overall milder phenotype. Finally, if the nucleotide sequence is the sole determinant of β-actin function, these mice would have no phenotype at all.

**Fig. 1.**
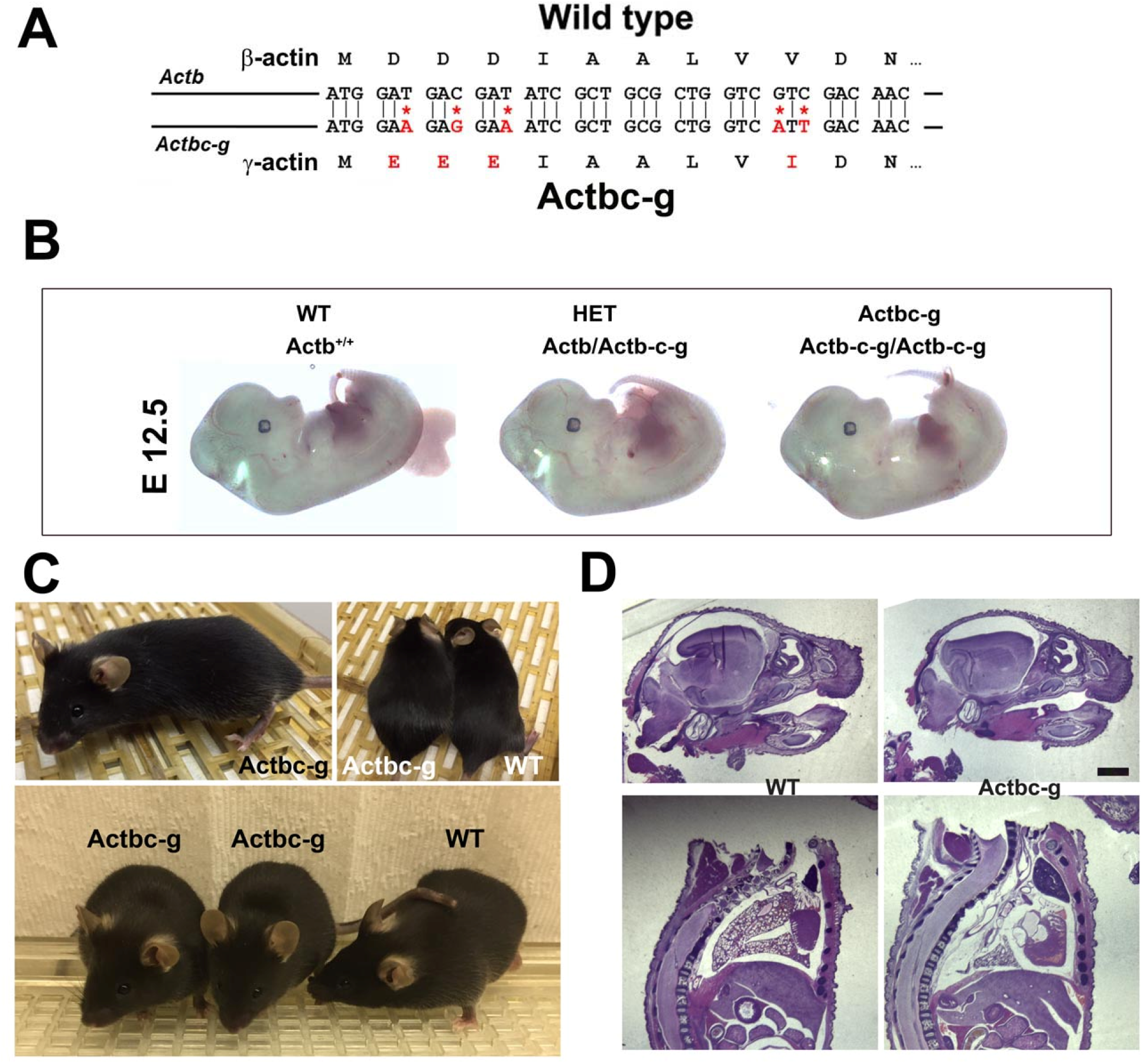
β -coded γ -actin (*Actbc-g*) mice exhibit no phenotypic changes compared to control. A, CRISPR/Cas9 editing strategy used to generate *Actbc-g* mouse. B, photos of *Actbc-g* E12.5 mouse embryos, with genotypes indicated. C, photos of *Actbc-g* mice after gene editing, alone (top left) and next to age-matched (top right) and littermate wild type (WT) (bottom). Three mice from two different litters are shown. D, H&E-stained sagittal sections of the heads (top) and bodies (bottom) of littermate P0 wild type (WT) and *Actbc-g* mice. Scale bar, 1 mm.

This gene editing strategy was successful, generating homozygous mouse mutants that contained no β-actin protein (Fig S1, S2, and S3). Strikingly, *Actbc-g* mice appeared completely healthy, viable, and fertile with no signs of deficiencies previously seen in any of the β-actin knockout mouse models. These mice did not exhibit any visible defects in embryogenesis (Fig. 1B), and appeared healthy and normal after birth (Fig. 1C) (observed, in one case, until approximately 8 months old). These mice also had normal fertility, as evidenced by litter sizes from *Actbc-g* homozygous breeding pairs that averaged 6.4 pups per litter (±0.38 SEM, n=9), compared to the average litter size of 6.3 pups previously reported for their matching background wild type strain C57BL/6 (http://www.informatics.jax.org/silver/tables/table4-1.shtml). Thus, this result definitively proves that β-actin nucleotide sequence, rather than its amino acid sequence, determines the essential function of β-actin *in vivo*.

To test for possible milder defects in these mice, we analyzed the overall morphology and appearance of all major organs and body parts in newborn (P0) *Actbc-g* mice by sagittal sectioning and H&E staining, and found no overall differences or abnormalities between wild type and *Actbc-g* mice (Fig. 1D), suggesting that the embryonic development in these mice occurs normally. Overall, *Actbc-g* mice appeared completely healthy and normal, suggesting that γ-actin protein encoded by the β-actin gene is fully able to functionally substitute β-actin’s essential role in mouse survival and health.

To confirm the replacement of β-actin protein in these mice with the γ-actin protein, we performed quantitative Western blots from several tissues where non-muscle actin isoforms are normally expressed at high levels, including brain, kidney, liver, and lungs (Fig. 2). In all these tissues, loss of β-actin protein was accompanied by a prominent increase in γ-actin, without overall changes in total actin levels (Fig. 2 and S3). Corresponding changes were also seen on 2D gels from these tissues, run under shallow pH gradient to separate actin isoforms (Fig. S4).

**Fig. 2.**
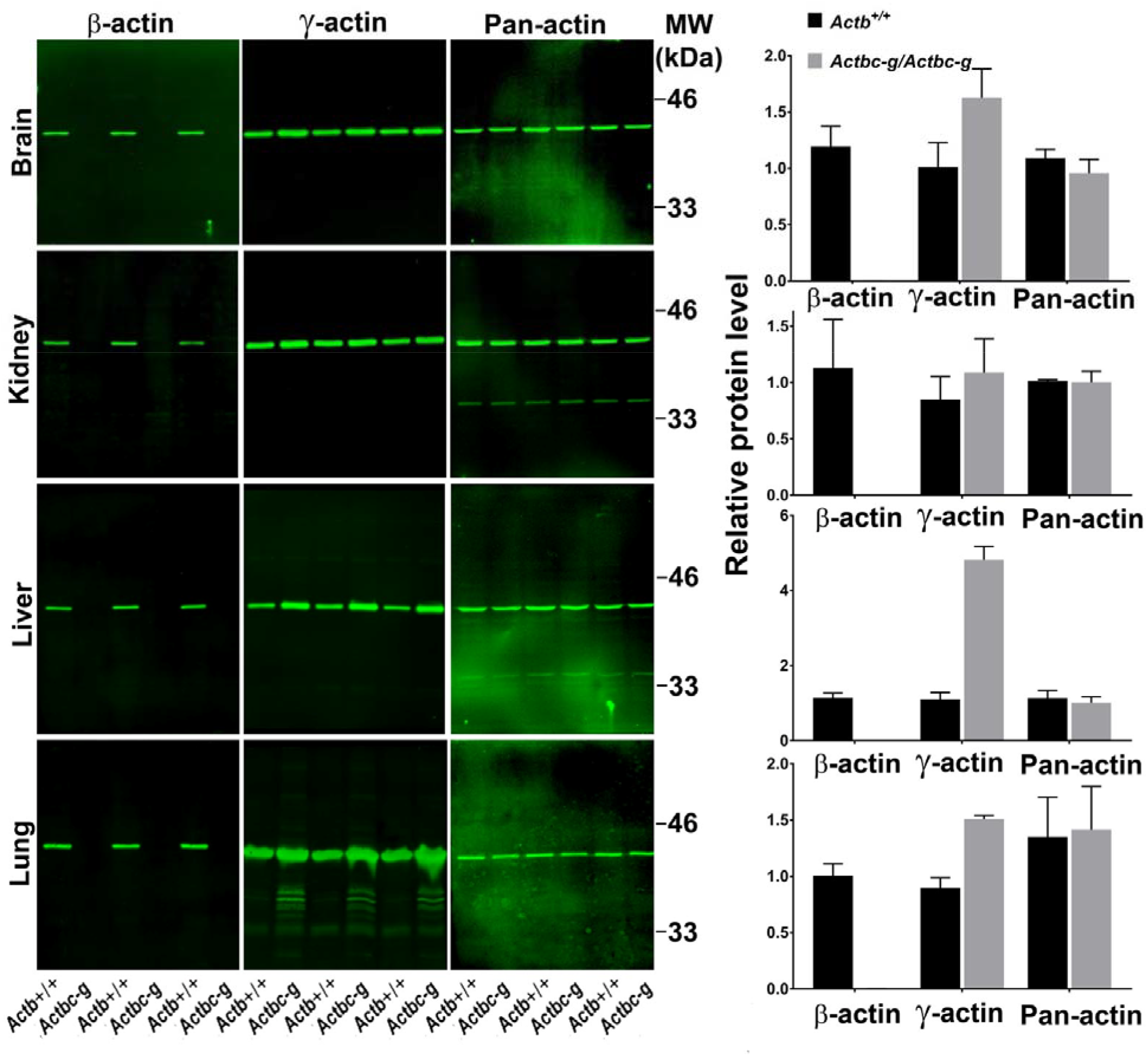
*Actb* gene editing abolishes β-actin protein from multiple organs and is accompanied by up-regulation of γ-actin without changing the total actin levels. Western blot analysis showing images (left) and quantifications (right) of whole tissue lysates from wild type (*Actb+/+*) and *Actbcg* mice. Fluorescence images obtained from the Odyssey gel imager are shown. For quantification, total fluorescence from the 43 kDa actin band was normalized to the loading control and to the actin level in the first lane for each blot. Error bars represent SEM, n=3.

We also performed a reciprocal experiment, using CRISPR/Cas9 gene editing to edit the mouse γ-actin gene to encode β-actin protein (“gamma-coded beta actin” or γc-β-actin, Fig. S5–S6). This strategy was only partially successful, resulting in replacement of the first three nucleotides to convert the N-terminal MEEE sequence of γ-actin into the MDDD sequence of β- actin, while failing to achieve the V/I substitution at codon 10. However, given that the full deletion of γ-actin has a much milder phenotype than β-actin mouse knockout (Belyantseva et al., 2009; Bunnell et al., 2011; Bunnell and Ervasti, 2010), we did not pursue this further and analyzed the partially edited mouse instead. The *Actg1c-b* mice showed disappearance of γ-actin protein and a corresponding increase in β-actin protein by Western blots (Fig S7). These results strongly suggest that γ-actin *in vivo* functions, like β-actin, is also defined by its nucleotide, rather than amino acid sequence.

### γ-actin protein expressed off β-actin gene supports normal cell migration

Since β-actin has been previously shown to play a major role in directional cell migration, and its knockout in cells leads to severe impairments in their actin cytoskeleton organization and their ability to migrate (Bunnell et al., 2011; Tondeleir et al., 2012), we next analyzed the actin cytoskeleton distribution and directional migration of mouse embryonic fibroblasts (MEF) derived from littermate wild type and *Actbc-g* mice. Despite the complete absence of β-actin protein in these cells, their actin cytoskeleton appeared similar to that of wild type cells. We detected no difference in F-actin levels in these cells (Fig. 3), or in the morphology and appearance of the actin cytoskeleton (Fig. 3, 4).

**Fig. 3.**
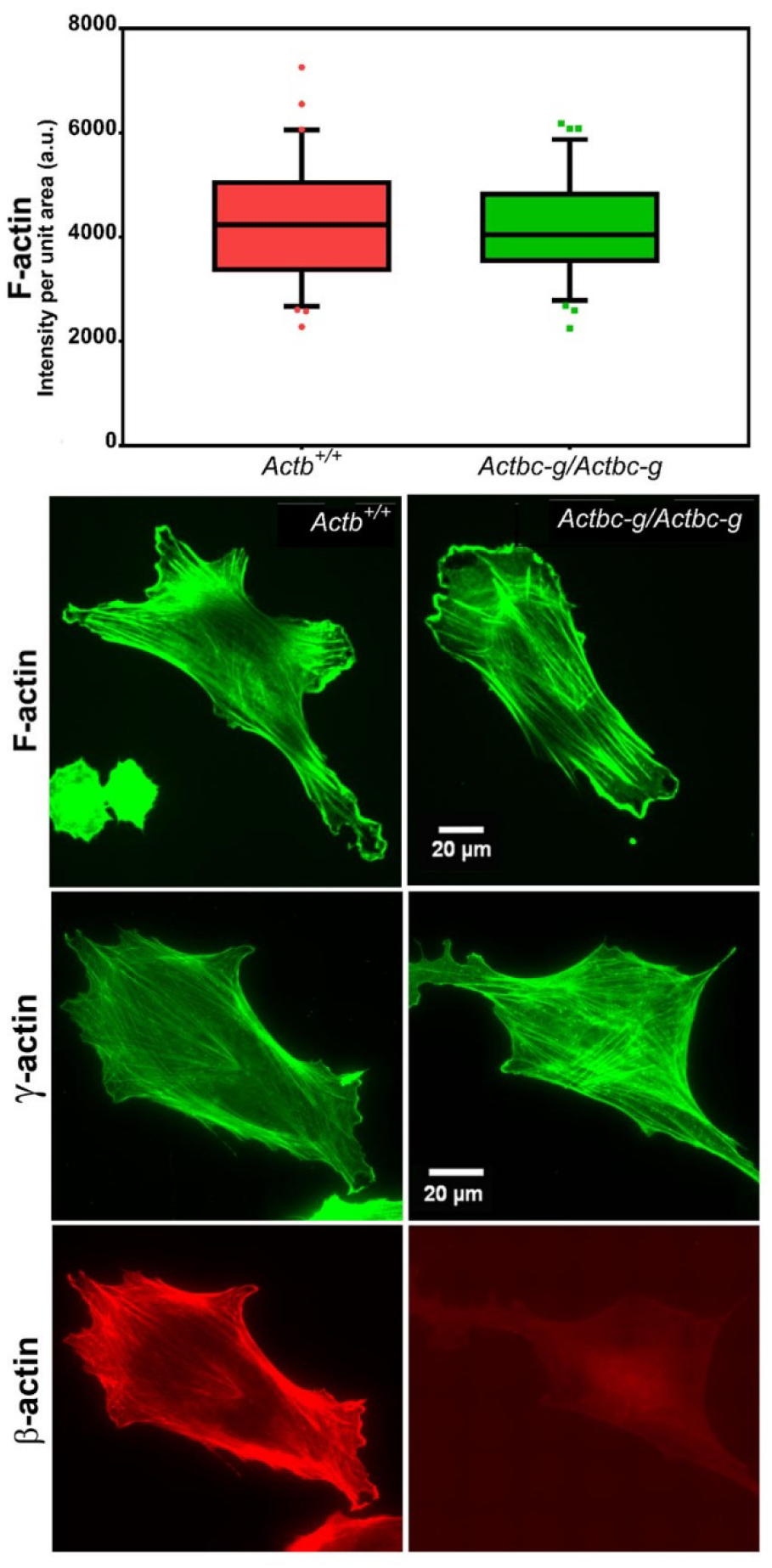
Mouse embryonic fibroblasts derived from *Actbc-g* mice have normal actin cytoskeleton, despite complete lack of β-actin. Top, quantification of total F-actin detected by Phalloidin-AlexaFluor488 staining of wild type (Actb+/+) and *Actbc-g* primary mouse embryonic fibroblasts. Numbers were averaged from 69 cells in WT and 76 cells in *Actbc-g*, obtained from 2 different primary cultures independently derived from 2 different littermate embryos for each set. Bottom, representative images of both cell types stained with Phalloidin-AlexaFluor488 or antibodies to both actin isoforms as indicated.

**Fig. 4.**
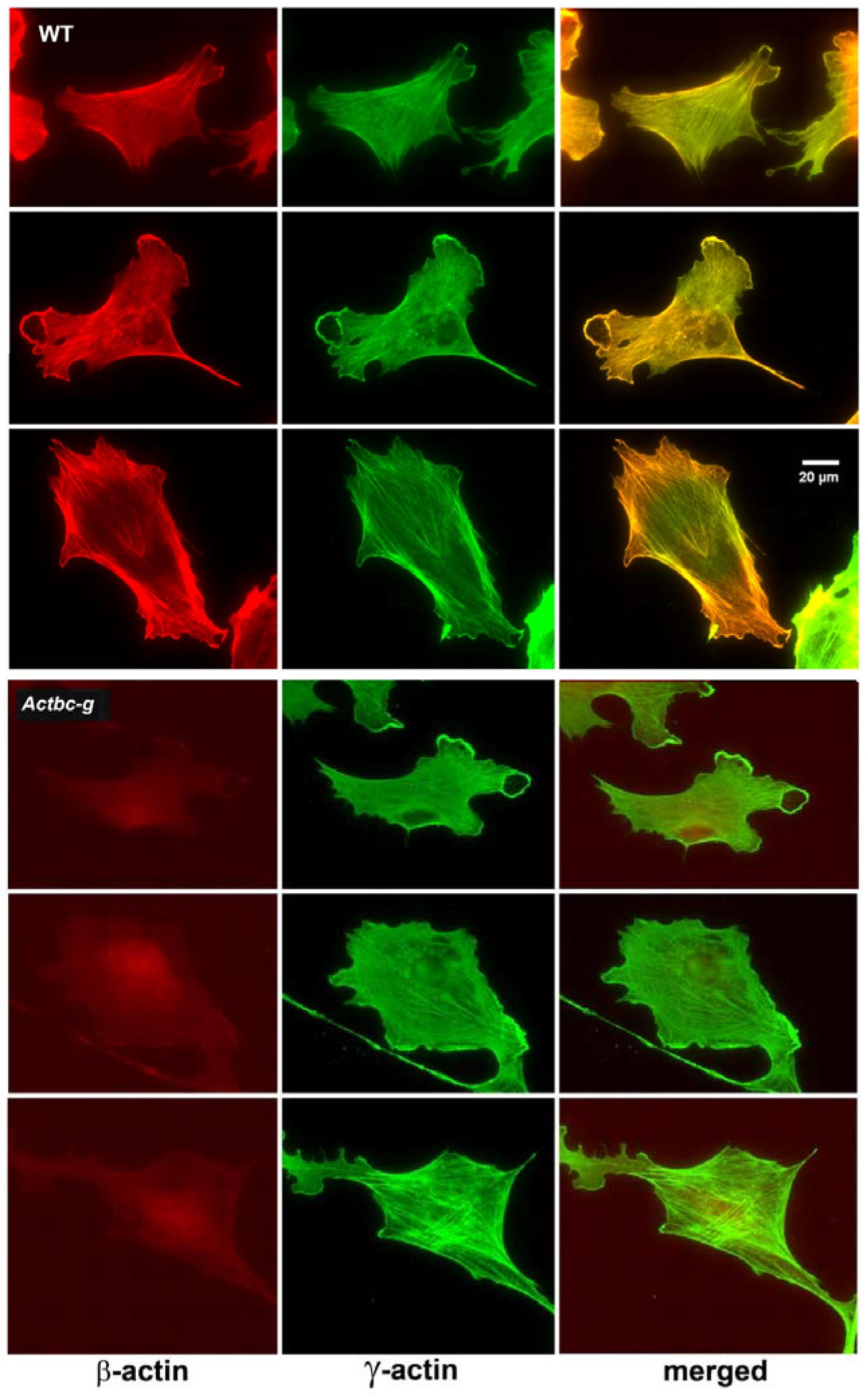
Mouse embryonic fibroblasts show no major changes in morphology and actin distribution. Representative images of wild type (WT) and *Actbc-g* primary mouse embryonic fibroblasts stained with antibodies to both actin isoforms as indicated.

To test the ability of these cells to migrate, we performed wound healing assays to measure the overall migration rates of the cell monolayers in wild type and *Actbc-g*. We also measured the directionality of single cell migration on fibronectin-coated dishes. In both assays, no difference was observed between the two cell types (Fig. 5), confirming that the actin isoform substitution did not result in any significant changes in these cells’ ability to migrate. Thus, our data definitively demonstrate that the essential function of β-actin *in vivo* is defined by its nucleotide, and not its amino acid sequence.

**Fig. 5.**
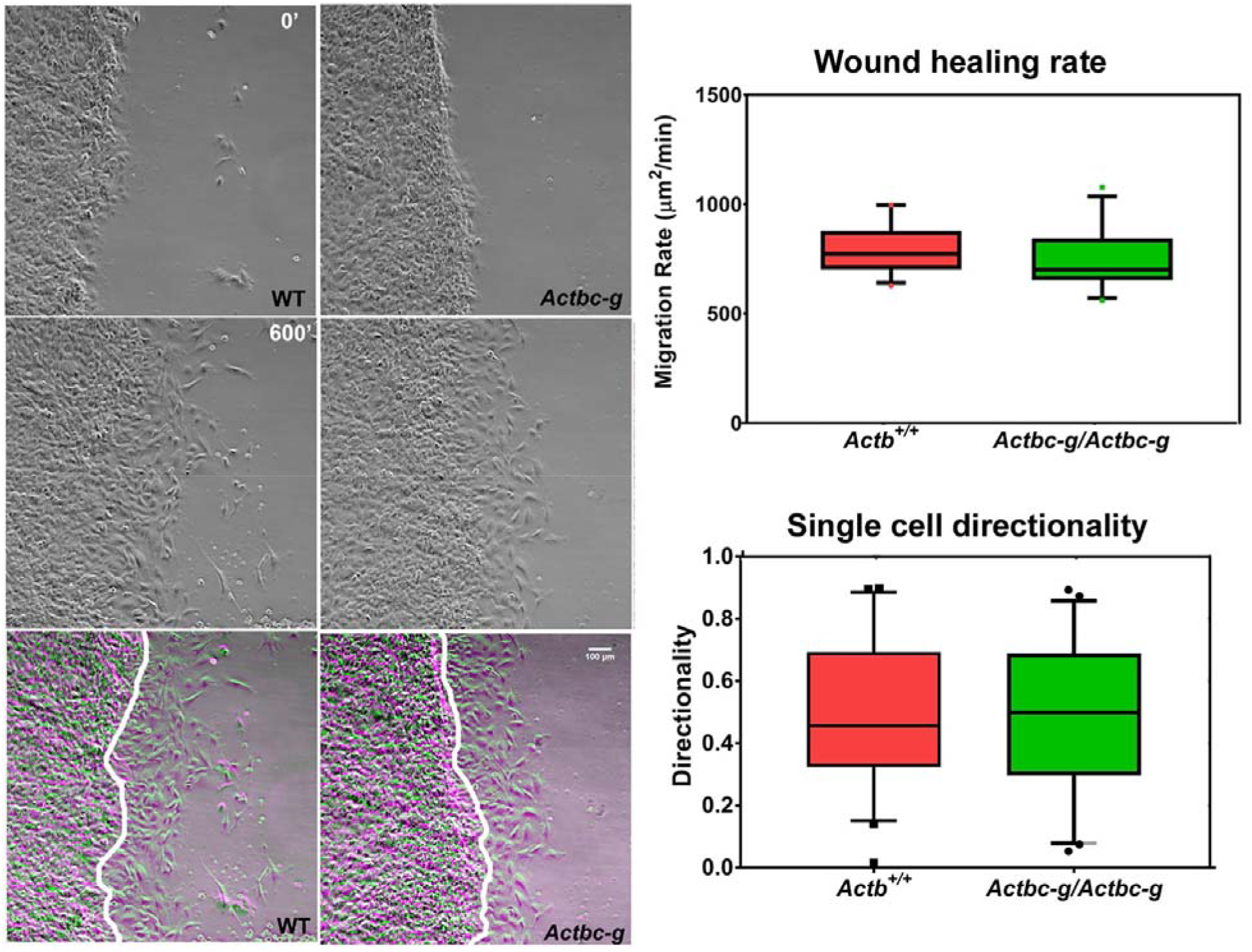
Mouse embryonic fibroblasts derived from *Actbc-g* mice migrate at normal rates. Left, phase contrast images of the first (0’) and last (600’) frame taken from a representative time lapse videos of the WT and *Actbc-g* cells at the edge of a monolayer migrating into an infinite scratch wound. Overlay of the two frames is shown in the bottom row. Scale bar, 100 μm. Right top, quantification of the cell migration rate as μm^2^/min, WT: n=28, *Actbc-g*. n=29 averaged from two independently derived primary cultures for each set. Right bottom, quantification of cell directionality in single cell migration assays (calculated as persistence over time; n=49 for WT and 50 for *Actbc-g*) suggests that single cell migration was not affected in *Actbc-g* cells.

## Discussion

Our results demonstrate a novel determinant – silent substitutions in nucleotide sequence – that drives the differential functions of non-muscle actin isoforms, a mechanism potentially applicable to other members of the actin family and other highly similar proteins in eukaryotic genomes. In the case of non-muscle actins, our finding resolves decades of controversial studies and reconciles a body of seemingly contradictory results obtained in the attempts to address functional distinction between β- and γ-actin (Bergeron et al., 2010; Dugina et al., 2009; Kapustina et al., 2016; Patrinostro et al., 2017). Our data demonstrate that the nucleotide sequence of the gene, rather than the amino acid sequence of the encoded isoform, determines the absolute requirement of β-actin for organism’s survival.

Closely related protein isoforms can exhibit functional differences which can be attributed to one or more of the following three sources. First, variations at the amino acid level can cause profound differences in protein function. Second, variations in mRNA properties due to differences in their coding sequence and UTR regions can strongly affect mRNA localization, stability, and translatability via secondary structure and codon usage. Finally, variations in gene intron sequences, promoter, and enhancer regions can contribute to the overall gene regulation, expression levels, and tissue specificity. Contribution of each of these levels to actin isoform function has been extensively investigated in prior studies. While β-actin and γ-actin isoforms share a remarkable conservation at the amino acid level, with just four homologous amino changes at their N-termini, these two proteins have been shown to have slightly different polymerization kinetics (Bergeron et al., 2010) and to differentially interact with cofilin (Kapustina et al., 2016) and non-muscle myosin isoforms (Muller et al., 2013). At the level of mRNA, actin 3′UTRs are isoform-specific and evolutionarily conserved, suggesting that they play important roles *in vivo* (Erba et al., 1986; Hill and Gunning, 1993). A plethora of literature elucidates the importance of β-actin 3′UTR for the localization of the transcript (see, e.g. (Condeelis and Singer, 2005) for a comprehensive review). It has also been shown that γ-actin mRNA induces a ribosome pausing event, resulting in its slower translation compared to β-actin, a mechanism that drives their differential arginylation (Zhang et al., 2010). An alternative poly A site in β-actin mRNA increases its translation levels (Ghosh et al., 2008). Finally, at the gene level, several studies point to roles of various regulatory elements in the actin isoforms. γ-actin gene contains a unique and highly conserved intron III (Lloyd and Gunning, 1993), and an alternatively spliced exon 3a (Drummond and Friderici, 2013). β-actin exhibits both 3UTR-dependent (Lloyd and Gunning, 1993; Lyubimova et al., 1999) and 3’UTR-independent (Lloyd et al., 1992; Schevzov et al., 1992) feedback regulation of gene expression.

Our result that targeted genome editing of mouse β-actin gene to encode for β-actin protein leads to no apparent phenotype proves for the first time that the major determinants of β-actin’s essential function *in vivo* are encoded at the nucleotide level. Some of the existing data point to the possibility that this effect is mediated primarily or exclusively via the coding region, rather than the promoter, the UTR, or the intron sequences. Indeed, deletion of exons 2 and 3 of the β- actin gene that include β-actin translation initiation site but do not encompass the promoter region, UTR, or the non-coding elements in the rest of the gene leads to embryonic lethality (Bunnell et al., 2011). Notably, in *Actb* knockout mice other actin isoforms are up-regulated to compensate for the total actin dosage, but this promoter-mediated up-regulation is insufficient to rescue the phenotype of early embryonic lethality. At the same time, targeted insertion of the human β-actin coding sequence into this region rescues embryonic lethality in these mice, further supporting the idea that the coding sequence plays a key role (Tondeleir et al., 2012). Data from our group previously showed that coding sequence drives differences in actin’s posttranslational modifications, one of the forms of functional actin regulation (Zhang et al., 2010). While these studies do not fully exclude a potential contribution of non-coding elements, especially those that may be located between exons 2 and 3 in the gene, they strongly support our hypothesis that the coding region is primarily responsible for the uniquely essential role of β-actin *in vivo*. Elucidating the underlying hierarchy of silent substitutions among other modes of regulation, would further our understating of the various levels of factors effecting the function of different actin isoforms.

## Methods

### CRISPR/Cas9 mutagenesis and genotyping

C57Bl/6 strain was used to generate the gene-editted mice. The donor females were superovulated using 5 IU of PMSG followed 48 hours later with 5 IU HCG, after which the females were mated immediately to C57Bl/6 studs. MEGAshort-script T7 transcription kit (Ambion) was used for in vitro transcription of small guide sgRNA (gctgcgctggtcgtcgacaaCGG, where CGG is the Protospacer Adjacent Motif, PAM) as per manufacturer’s protocol. mMESSAGE mMACHINE T7 transcription kit (Ambion) was used to synthesize Cas9 mRNA. MEGAclear transcription clean up kit (Ambion) was used to purify the synthesized RNAs. About 20 hours post HCG, the CRISPR solution was injected into zygotes via pronuclear injection at a concertation of: Cas9 mRNA: 100ng/μL, template DNA: 100ng/μL, and gRNA: 50ng/μL. The zygotes were further cultured overnight in KSOM media using a 5% CO2 incubator. All the embryos which successfully cleaved to the 2-cell stage were transferred into recipient females via oviduct transfer. A founder female that was mosaic for the mutation was derived and crossed with a wildtype male to derive heterozygotes. One male and female heterozygote from the F1 generation were crossed to produce F2 generation. Two separate litters from the F2 generation produced 2 wildtype females, 5 heterozygote males, 1 heterozygote female, and 3 homozygote males.

Template DNA sequence:

5’CGGCTGTTGGCGGCCCCGAGGTGACTATAGCCTTCTTTTGTGTCTTGATAGTAGTTCGCCATGGAAGAGGAAATCGCTGCGCTGGTCATTGACAACGGCTCCGGCATGTGCAAAGCCGGCTTCGCGGGCGACGATGCTCCCCGGGCTGTA 3’

The *Actb* gene has an EcoRV site which gets destroyed upon gene editing (Fig. S1). We utilized restriction digestion of a PCR product produced from the *Actb* gene to determine the genotype of the resulting mice. While wild type (Actb^+/+^) mice gave two bands: 600bp and 300bp upon EcoRV digestion, mice homozygous for the mutations (*Actbc-g/Actbc-g*) give a single band at 900bp. PCR products from mice that are heterozygous for the mutation (*Actb/Actbc-g*) gave three bands upon EcoRV digestion (Fig. S1). The results were further verified by sequencing the 5’ end of the *Actb* gene. In order to verify that β-actin protein was no longer produced, tail samples were lysed and a western blot was carried out using isoform specific antibodies: mouse anti-β-actin (Clone 4C2, EMD Millipore and Clone AC15, Sigma Aldrich), mouse anti-γ-actin (Clone 2C3 EMD Millipore).

### Cell culture

Primary Mouse Embryonic Fibroblasts (MEFs) were isolated from the back area of freshly euthanized E12.5 mouse embryos by tissue disruption and cultured in DMEM (Gibco) supplemented with 10% FBS (Gibco).

### Cell migration assays and imaging

Cell migration was stimulated by making an infinite scratch wound. The cells were allowed to recover for a period of 2 hours before imaging. Migration rates were measured as the area covered by the edge of the wound in the field of view per unit time using Fiji (NIH). For measurements of directionality in single cell migration assays, primary MEFs were cultured on glass bottom MatTek dishes coated with 5ug/ml Fibronectin at low cell densities. 2 hours after seeding, cells were imaged at 10 minute intervals for 4 hours. Single cells were tracked using Metamorph (Molecular Devices) Track Objects module. The obtained total displacement was divided by total distance of the track to obtain a directionality score shown in Fig. 5. All images for these experiments were acquired on a Nikon Ti microscope with a 10X Phase objective and Andor iXon Ultra 888 EMCCD camera.

### Immunofluorescence

To quantitfy the amount of actin polymer, cells were seeded on coverslips in six well plate at 20, 000 cells/well overnight and fixed in 4% (w/v) PFA at room temperature for 30 minutes. Cells were then stained with Phalloidin conjugated to AlexaFluor 594 (Molecular probes). Images were acquired using Andor iXon Ultra 888 EMCCD camera at 40X and the total intensity of phalloidin per cell was measured using Metamorph (Molecular Devices).

### Western blotting

Tissues from 2 month old *Actb^+/+^* and *Actbc-g/Actbc-g* mice were collected and flash frozen in liquid nitrogen. Brain, Kidney, Liver and Lung tissues were ground and weighed. The samples were lysed directly in 4x SDS sample buffer (1:4 w/v). Equal volumes of the lysates loaded for SDS-PAGE and 2D gel electrophoresis. Following transfer of the gels, the blots were dried and stained with LI-COR REVRT Total protein stain as per manufacturer’s protocol. Images were obtained using an Odyssey scan bed in the 700nm channel. The blots were then blocked and incubated with primary antibodies for mouse anti-β-actin (Clone 4C2, EMD Millipore), mouse anti-γ-actin (Clone 2C3, EMD Millipore), and rabbit anti-pan-actin (Cytoskeleton, Inc.). Secondary antibodies against mouse and rabbit conjugated to IRDye800 were used to probe the blots and images were acquired in the 800nm channel using Odyssey scan bed. The total protein intensity was used to account for loading differences and the obtained signals were normalized to the first lane in the blot.

2D gel analysis was performed by Kendrick Laboratories, Inc. (Madison, WI) as described by (Burgess-Cassler et al., 1989; O’Farrell, 1975) using shallow pH gradient for the first dimension to separate actin isoforms (pH 4-6, 4-8).

## Acknowledgements

We thank Dr. John Pehrson for helpful discussions throughout this project and Drs. John Pehrson and Kei Miyamoto for critical reading of the manuscript. This work was supported by NIH grants GM104003 and GM117984.

## Supplementary Materials

**Fig. S1.**
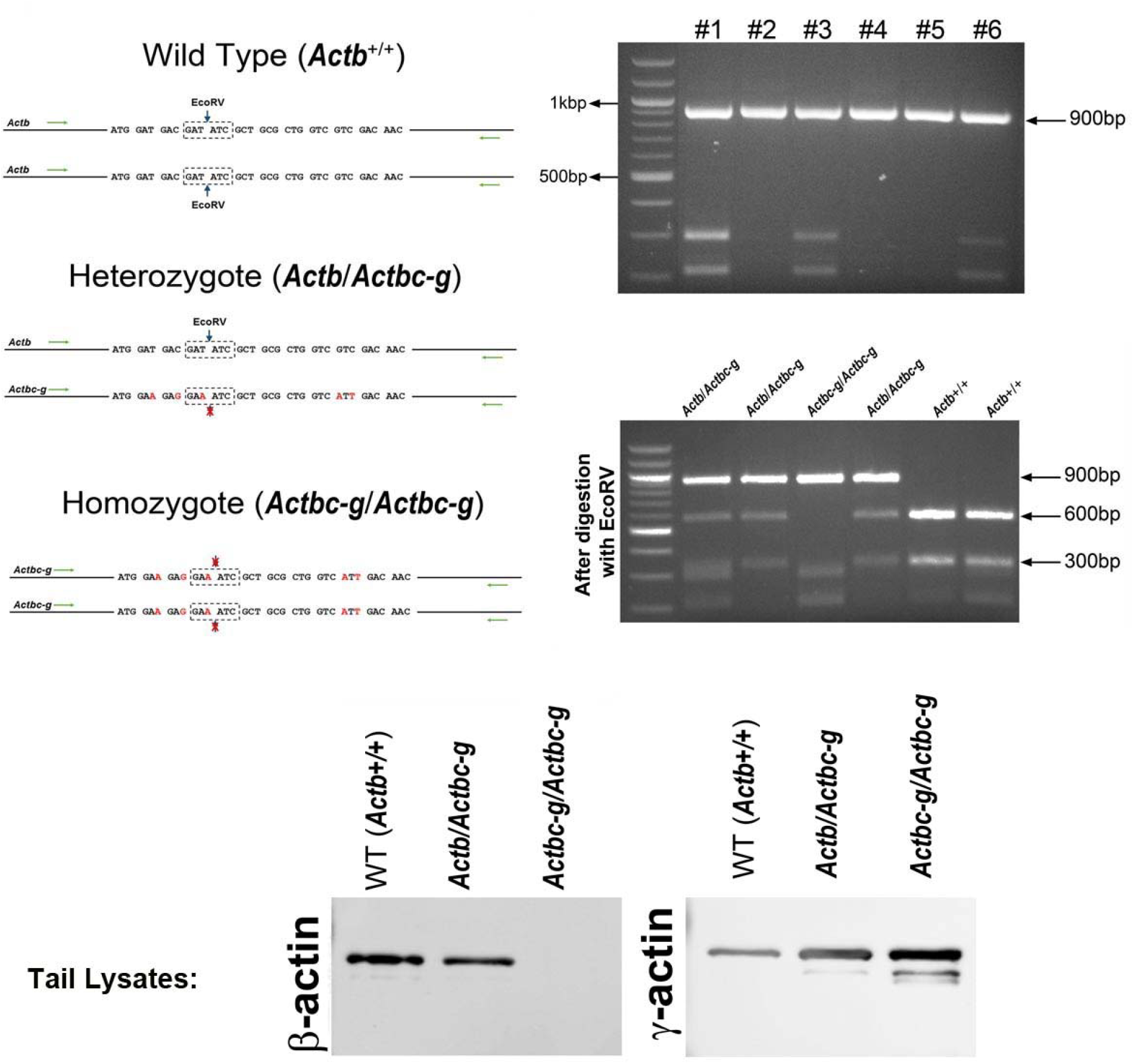
Generation of *Actbc-g* mouse. Top left, genotyping strategy: editing of the N-terminal codons of the β-actin gene abolishes an EcoRV restriction site, enabling the screening of the edited gene variants by EcoRV digestion of the PCR-generated DNA fragments corresponding to the 5’ of the actin sequence. Top right, PCR products before (top) and after (bottom) EcoRV digestion. Bottom, Western blots of wild type, heterozygous, and *Actbc-g* mouse tail lysates with antibodies to β- and γ- actin.

**Fig. S2.**
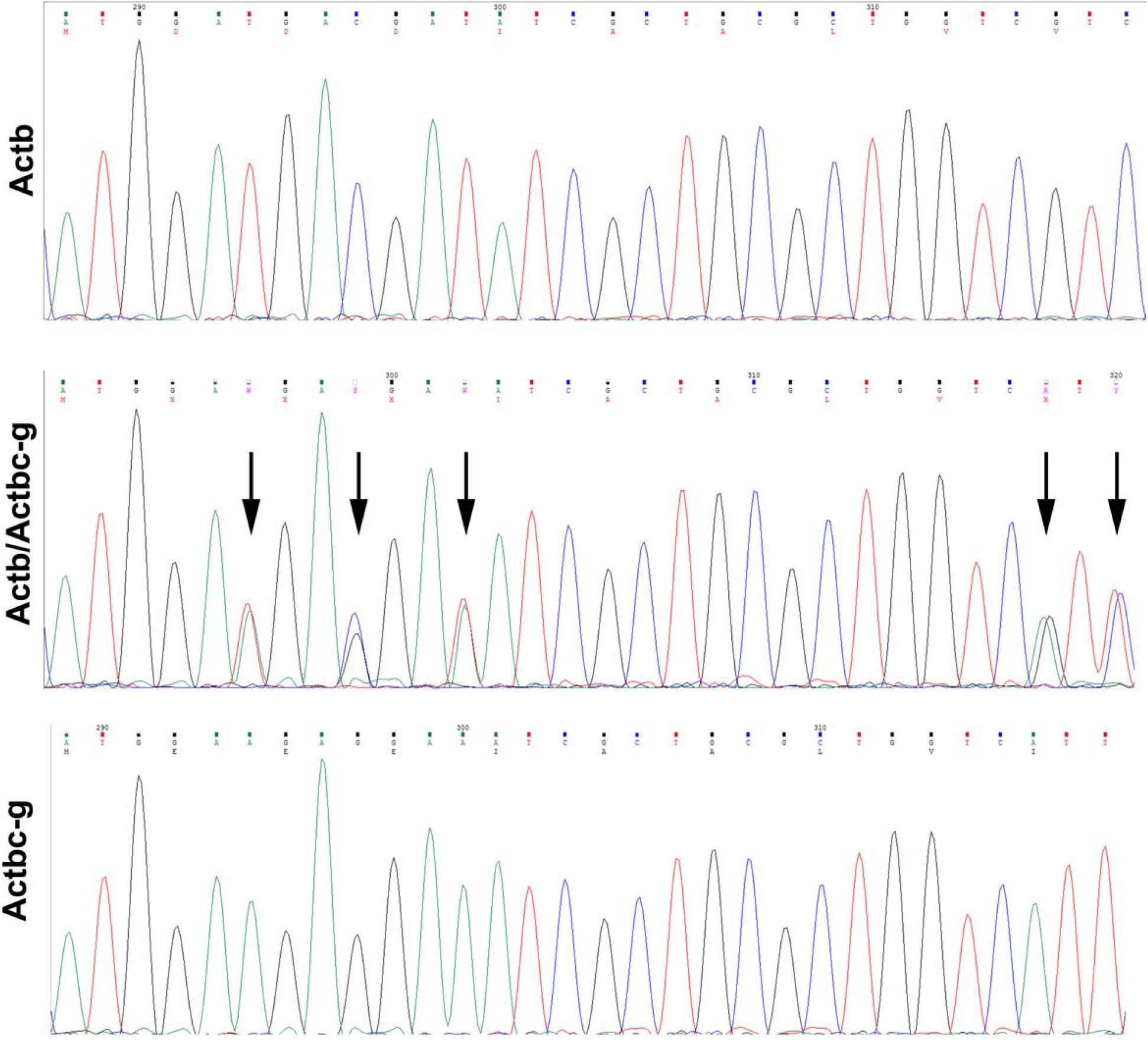
Sequencing result for wild type *Actb* and the edited *Actbc-g* alleles. Screen shots from the Chromas sequence viewing software.

**Fig. S3.**
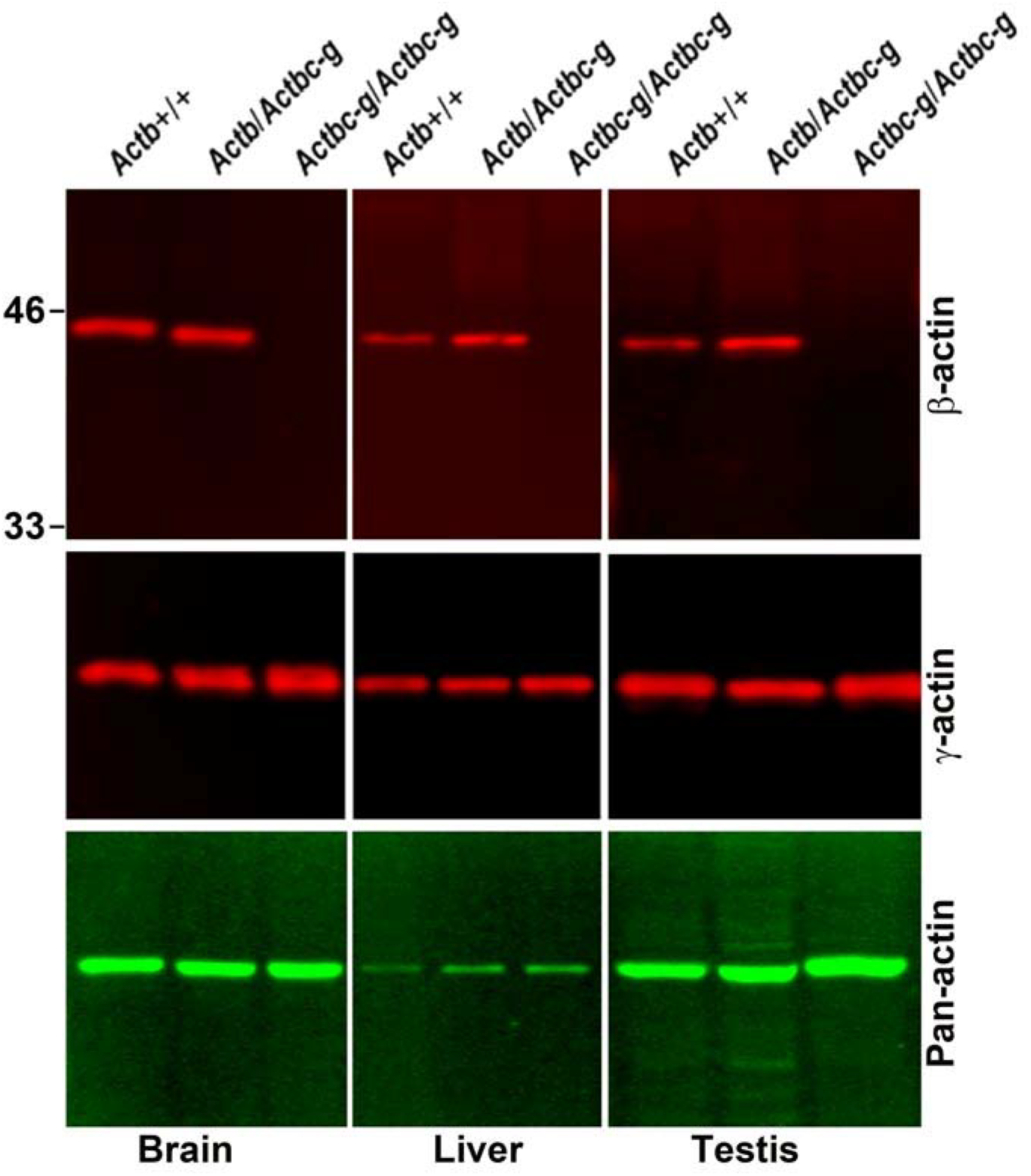
*Actb* gene editing abolishes β-actin protein from multiple organs and is accompanied by up-regulation of γ-actin without changing the total actin levels. Western blots of wild type, heterozygous, and *Actbc-g* mouse brain lysates probed with antibodies to β- and β-actin and total actin (pan-actin). Mouse genotypes are indicated on top of each lane, and the antibodies used are listed below each blot.

**Fig. S3.**
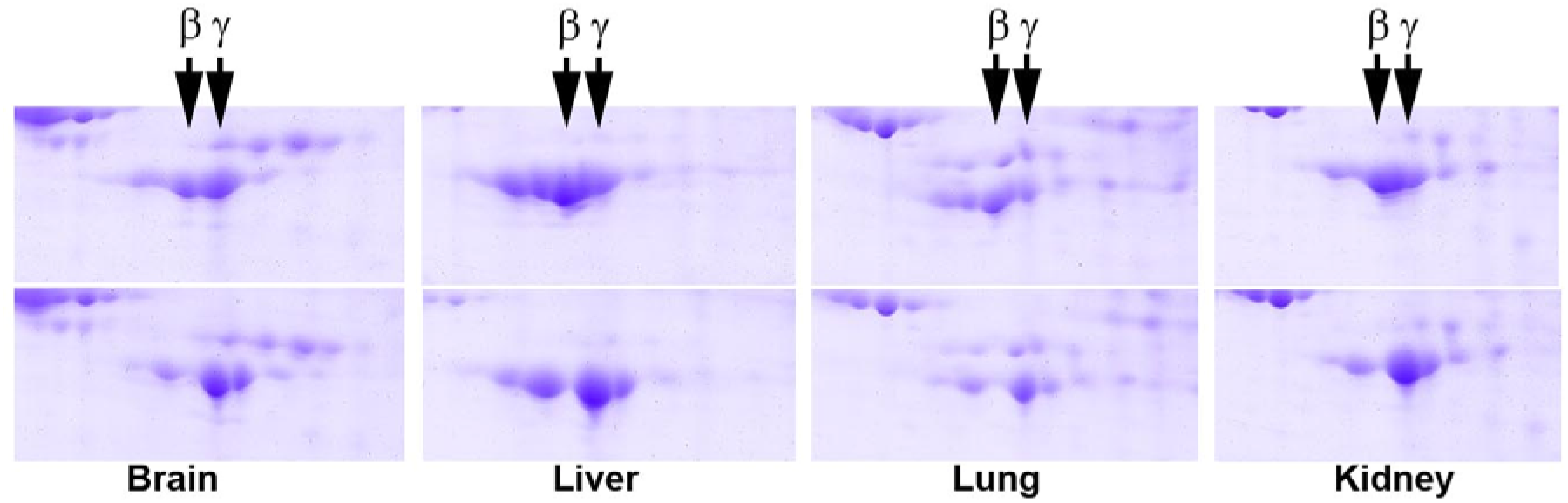
2D gel distribution of actin isoforms in wild type (top) and *Actbc-g* (bottom) mouse tissue lysates. Arrows indicate the position of the main spots for the two non-muscle actin isoforms, as labeled.

**Figure S5.**
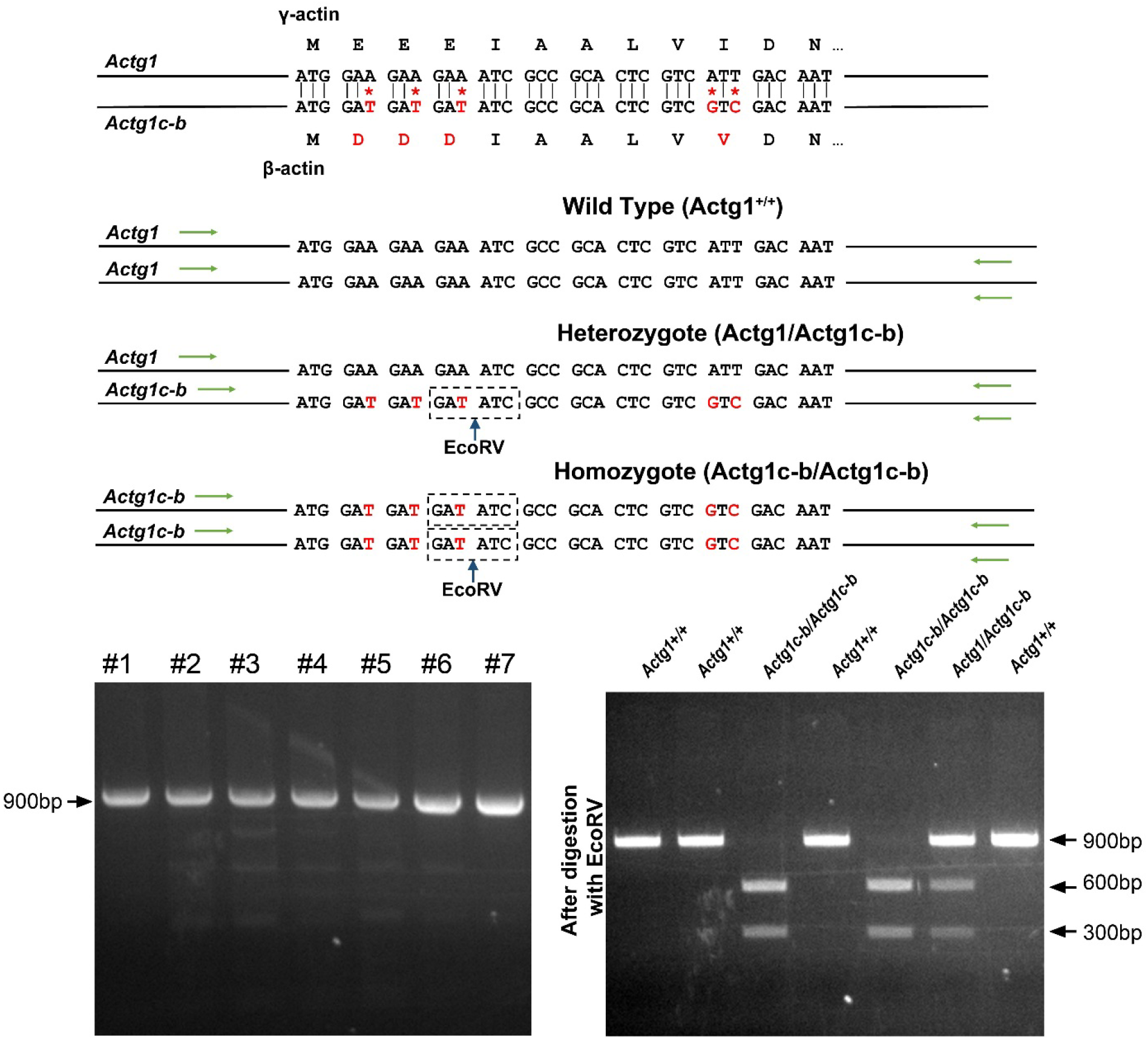
Generation of *Actg1c-b* mouse. Top, genotyping strategy: editing of the N-terminal codons of the γ- actin gene generates an EcoRV restriction site, enabling the screening of the edited gene variants by EcoRV digestion of the PCR-generated DNA fragments corresponding to the beginning of the actin sequence. Bottom, PCR products before (top) and after (bottom) EcoRV digestion.

**Fig. S6.**
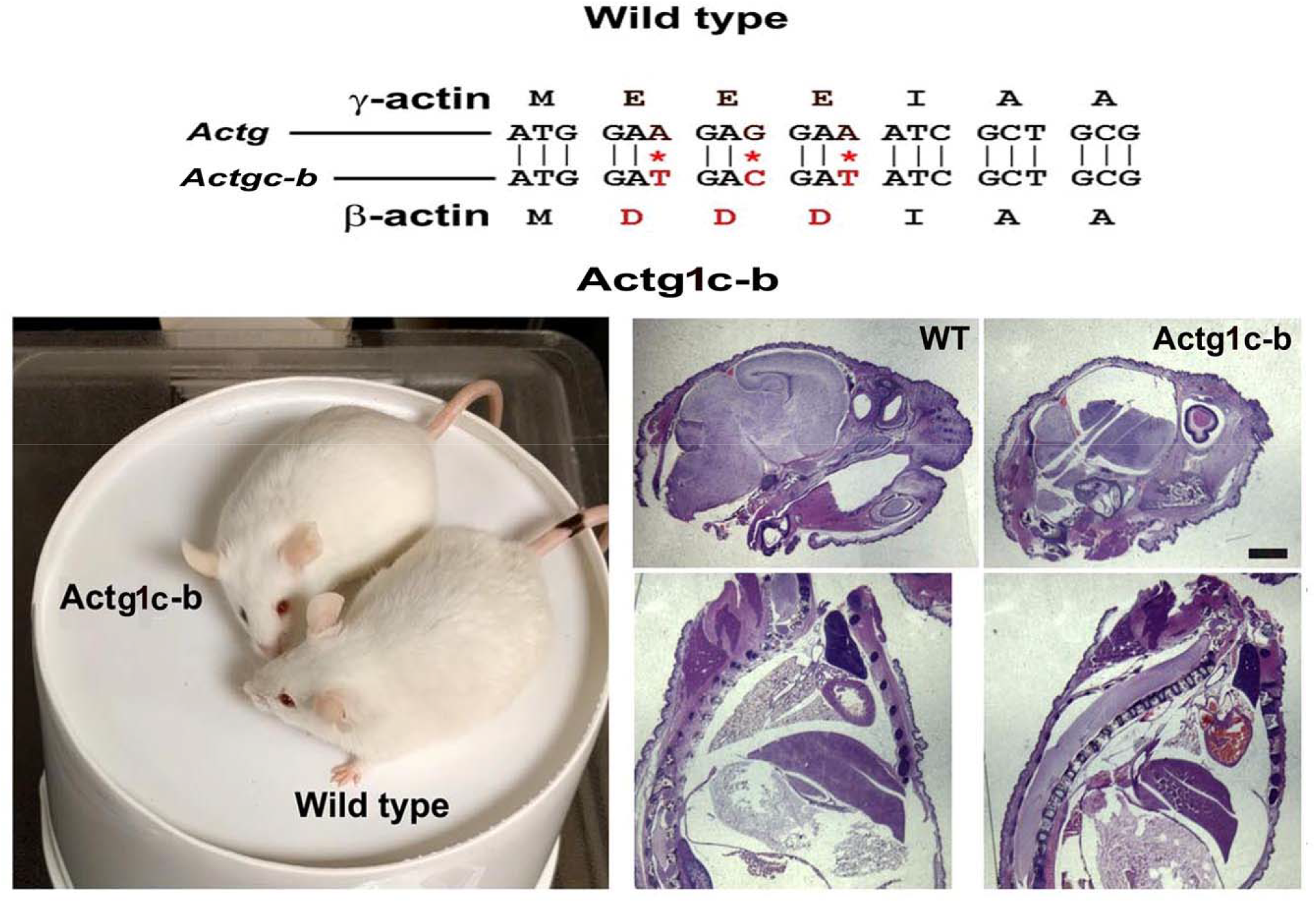
γ-coded β-actin (*Actg1c-b* mice exhibit no phenotypic changes compared to control. Top, CRISPR/Cas9 editing strategy used to generate *Actg1c-b* mouse. Bottom left, photos of *Actg1c-b* mouse after gene editing, to age-matched wild type. Bottom right, H&E-stained sagittal sections of the heads (top) and bodies (bottom) of littermate wild type (WT) and *Actg1c-b* mice. Scale bar, 1 mm.

**Figure S7.**
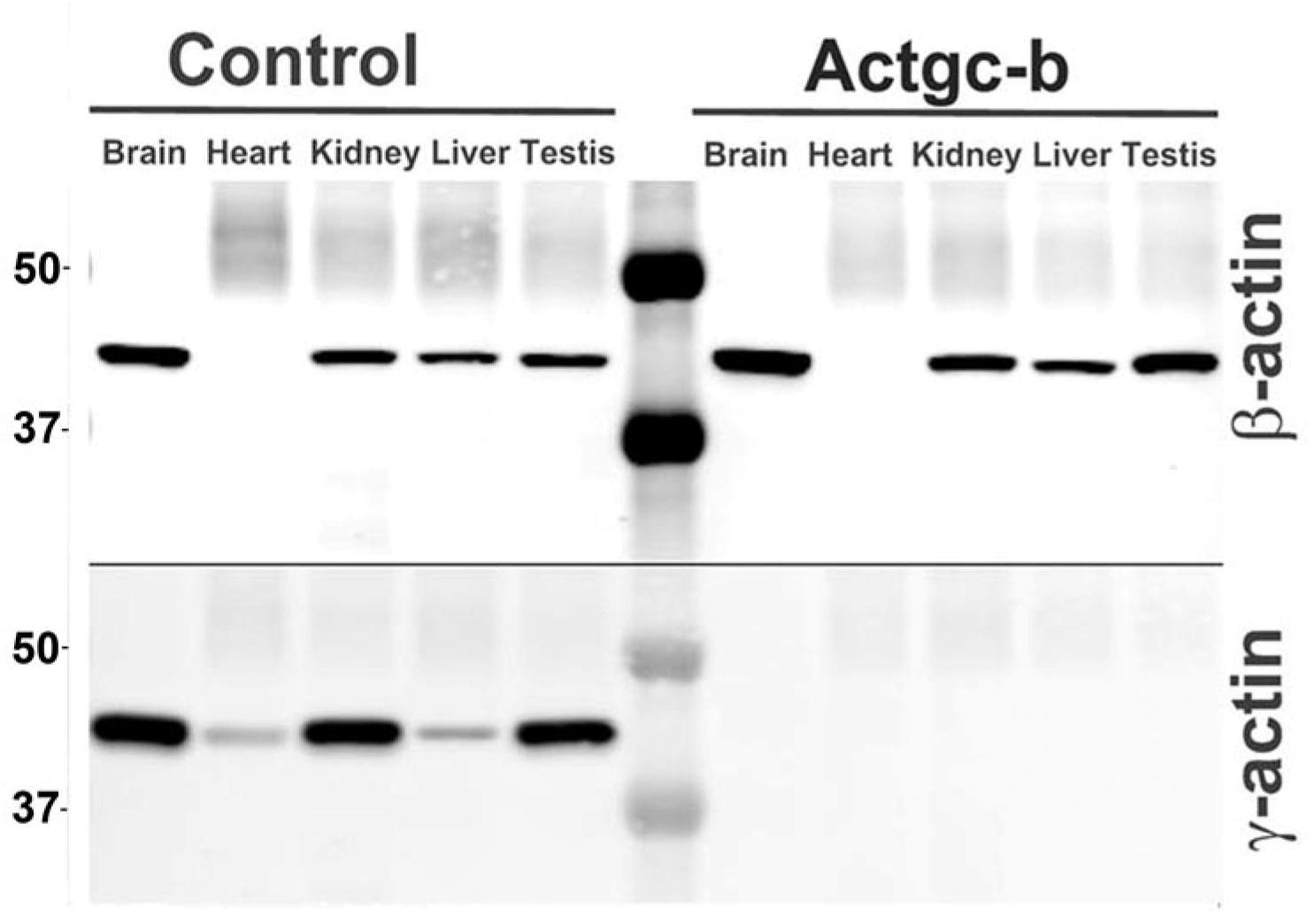
Partial editing of the γ-actin gene to encode β-actin-like protein abolishes γ- actin protein from multiple organs. Western blots with the actin antibodies indicated on the left using tissue homogenates from wild type control and *Actg1c-b* mouse.

